# A single test approach for accurate and sensitive detection and taxonomic characterization of Trypanosomes by comprehensive analysis of ITS1 amplicons

**DOI:** 10.1101/418343

**Authors:** Alex Gaithuma, Junya Yamagishi, Axel Martinelli, Kyoko Hayashida, Naoko Kawai, Megasari Marsela, Chihiro Sugimoto

## Abstract

The World Health Organization has targeted stopping the transmission of Human African Trypanosomiasis by 2030. To achieve this, better tools are urgently required to identify and monitor Trypanosome infections in human, animals, and tsetse fly vectors. This study presents a single test approach for detection and identification of Trypanosomes and their comprehensive characterization at species and sub-group level. Our method uses newly designed ITS1 PCR primers (a widely used method for detection of African Trypanosomes, amplifying the ITS1 region of ribosomal RNA genes) coupled to Illumina sequencing of the amplicon. The protocol is based on the widely used Illumina’s 16s bacterial metagenomic analysis procedure that makes use of multiplex PCR and dual indexing. We analyzed wild tsetse flies collected from Zambia and Zimbabwe. Our results show that the traditional method for Trypanosome species detection based on band size comparisons on a gel is unable to distinguish between *T. vivax* and *T. godfreyi* accurately. Additionally, this approach shows increased sensitivity of detection at species level. Through phylogenetic analysis, we identified Trypanosomes at species and sub-group level without the need for any additional tests. Our results show *T. congolense* Kilifi sub-group is more closely related to *T. simiae* than to other *T. congolense* sub-groups. This agrees with previous studies using satellite DNA and 18s RNA analysis. While current classification does not list any sub-groups for *T. vivax* and *T. godfreyi*, we observed distinct subgroups for these species. Interestingly, sequences matching *T. congolense* Tsavo (now classified as *T. simiae* Tsavo) clusters distinctly from the rest of the *T. simiae* Tsavo sequences suggesting that the Nannomonas group is more divergent than currently thought thus the need for a better classification criteria. This approach has the potential for refining classification of Trypanosomes and provide detailed molecular epidemiology information useful for surveillance and transmission control efforts.

**Author summary:** Detection of Trypanosomes in the tsetse flies plays an important role in the control of African trypanosomiasis by providing information on circulating Trypanosome species in a given area. We have developed a method that combines multiplex PCR and next-generation sequencing for Trypanosome species detection. The method is based on the widely used bacterial metagenomic analysis protocol and uses a modular, two-step PCR process followed by sequencing of all amplicons in a single run, making sequencing of amplicons more efficient and cost-effective when dealing with large sample sizes. As part of this approach, we designed novel primers for amplifying the ITS1 region of the Trypanosome rRNA gene that is more sensitive than conventional primers. Identification of Trypanosome species is based on BLAST searches against the constantly updated NCBI’s *nt* database, which facilitates the identification of Trypanosome subgroups. Our approach is more accurate than traditional gel-based analysis and shows how the latter is prone to misidentification. It is also sensitive and is able to discriminate between subgroups within Trypanosome species. Applied as an epidemiological tool, it has the potential to provide new, comprehensive and more accurate information on vector-pathogen-host interconnections which are key in the control and management of African trypanosomiasis.

## Introduction

Human African trypanosomiasis or sleeping sickness is classified as a neglected tropical disease by WHO, that is endemic in sub-Sahara Africa. Human African trypanosomiasis affects impoverished rural areas of sub-Saharan Africa, where it coexists with animal trypanosomiasis constituting a major health and economic burden [1]. The disease is caused by protozoan parasites of the genus *Trypanosoma*, it is transmitted by the bite of blood-sucking tsetse flies (Diptera, genus *Glossina*). The human disease is caused by *Trypanosoma brucei rhodesiense* and *Trypanosoma brucei gambiense*, causing an acute and chronic disease in humans respectively [2]. *T.b. rhodesiense* is found in East Africa and transmitted by *Glossina morsitans*, while *T.b gambiense* is distributed in West Africa and is mainly transmitted by *Glossina pallidipes* [3–5]. Uganda is the only country that both forms of the disease occur with the potential for overlapping infections [6]. The incidence of sleeping sickness has over the years, from 26,000 cases reported in 2000 to less than 8,000 cases reported in 2012 [7]. This decrease is attributed to improved case detection and treatment and vector management [8]. Despite this decreased incidence, it is estimated that up to 70 million people distributed over 1.5 million km^2^ remain at risk of contracting the disease [9]. Besides, African animal trypanosomiasis (AAT) is one of the biggest constraints to livestock production and a threat to food security in sub-Saharan Africa. The parasites *T. congolense* (Savannah) and *T. vivax* are considered the most important animal Trypanosomes due to their predominant distribution in sub-Saharan Africa and their economic impact due to their predominant distribution in sub-Saharan Africa and their economic impact [10]. They cause pathogenic infections in cattle (*Nagana*) and also infect sheep, goats, pigs, horses, and dogs. While *T. brucei brucei* (and *T. brucei rhodesiense*) is pathogenic to camels, horses, and dogs, but causes mild or no clinical disease cattle, sheep, goats and pigs. *T. simiae* causes a fatal disease in pigs and mild disease in sheep and goats. *T. godfreyi* shows a chronic, occasionally fatal disease in pigs experimentally [11,12]. *T. evansi* was originally found to infect camels but it is present in dromedaries, horses, and other equines as well as in a wide range of animals causing *Surra* disease, while *T. equiperdum* causes dourine in equines [13]. The latter two species are independent of the tsetse fly vector [14,15]. They are either transmitted mechanically for *T. evansi* or sexually for *T. equiperdum* therefore distributed outside sub-Saharan Africa. Given that Trypanosome parasites are maintained in wild and domestic animals as reservoirs, this complicates control and measures.

The ribosomal RNA (rRNA) sequence region harboring internal transcribed spacer (ITS) sequences have been used to identify Trypanosome species in hosts and vectors. Epidemiological and screening studies rely on PCR to amplify the internal transcribed spacer 1 (ITS1) region of ribosomal genes to analyze Trypanosome species diversity [16–19]. This locus located between the 18s and 5.8s ribosomal subunit genes with between 100–200 copies [16] and is widely used to identify Trypanosome species based on amplicon size in a gel. However, identification of *T.b. rhodesiense, T.b. gambiense, T.b. brucei* or *T. evansi*, specific detection is required. A major problem with ITS1 PCR besides sensitivity limitations compared to nested PCR, is the fact that widely used primers for ITS1 PCR amplification have major limitations in their detection capacity showing bias in detection of some Trypanosome species over others [17,18]. Some are prone to non-specific amplification particularly in bovine blood samples [19]. When dealing with a large number of samples either for tsetse fly or animal infection prevalence studies, undertaking multiple species-specific PCR for each sample is an expensive and a laborious undertaking. Most often it is preferred to sequence the ITS1 PCR amplicons to confirm species identification in favor of multiple PCRs, usually by capillary sequencing. Although next-generation sequencing (NGS) is a well-established method for profiling bacterial communities, with the exception of *Plasmodium* in mosquitoes, relatively few studies have applied this technology in the diagnostics of protozoal infections [20,21]. Next-generation sequencing allows high-throughput parallelization of sequencing reactions, is more sensitive and accurate at single nucleotide resolution (due to deep sequencing) and is therefore helpful to accurately determine the prevalence and genetic diversity of Trypanosome species in wildlife communities and potential vectors.

## Materials and methods

### Sample collection and extraction of DNA

Tsetse flies were obtained from Zambia; along the Kafue national park border (n=85, collected in June 2017) and from Rufunsa area (n=200) near Lower Zambezi National park (surrounding farms and villages) collected earlier in November and December 2013) (Fig 1). We also included 188 tsetse flies samples earlier collected from Hurungwe Game reserve in Zimbabwe between March and April 2014 to expand Trypanosome species spectrum. All flies collected in this study were caught on public land using Epsilon or customized mobile traps and preserved in silica gel. The dried flies were transferred to a smashing machine and crushed at 3,000 rpm for 45 sec. DNA was isolated using the DNA Isolation kit for mammalian blood (Roche USA) as per the manufacturer’s protocol with slight modification where solution I (Red blood cell lysis buffer) was not used. The DNA sample was stored at −80ºC until polymerase chain reaction (PCR).

**Fig 1.**
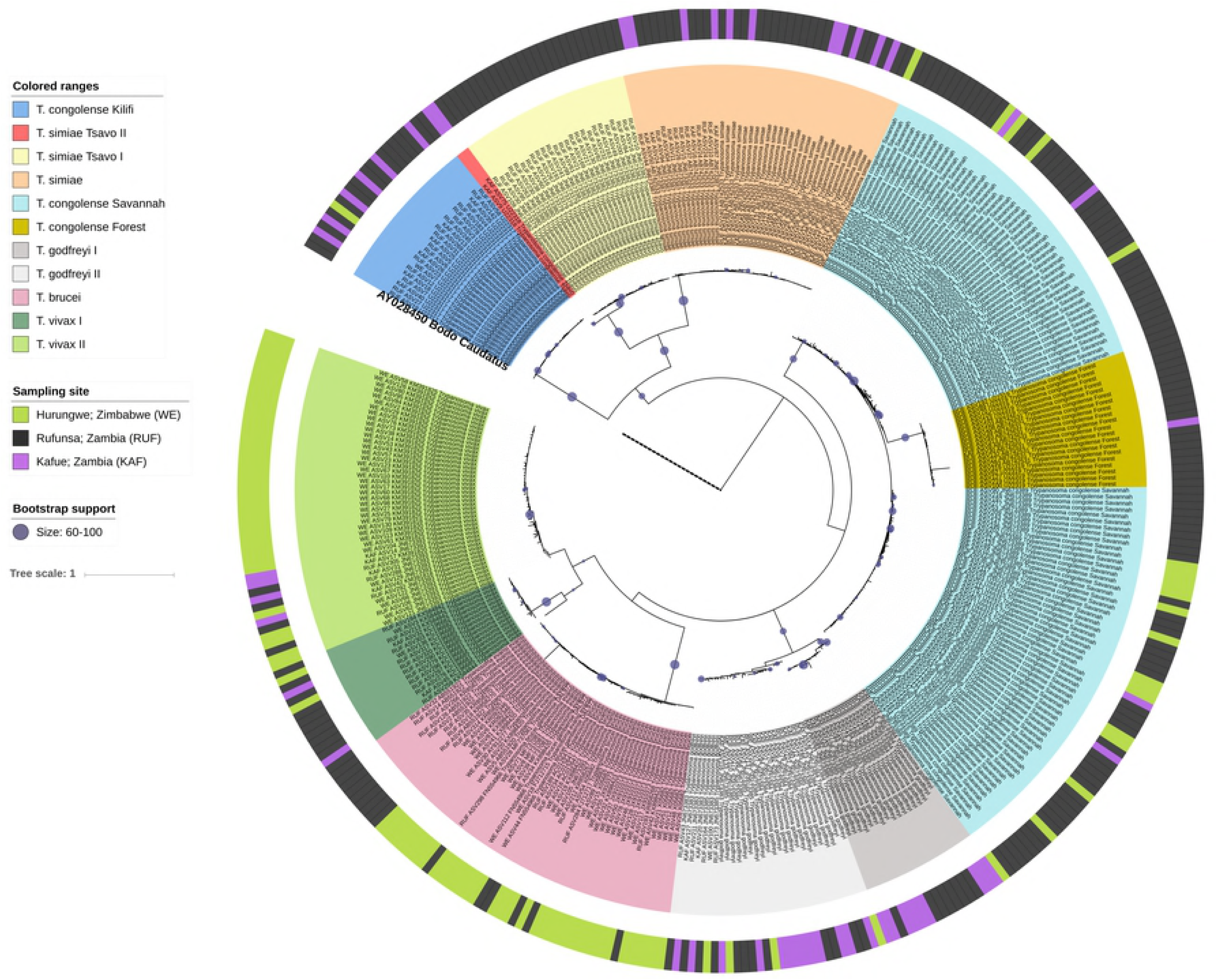
Map of Zambia and Zimbabwe showing areas of tsetse fly collection. Maps sourced from © OpenStreetMap contributors, and made available here under the Open Database License (ODbL) (https://opendatacommons.org/licenses/odbl/1.0/).

### Primer design and testing

The following sequences were retrieved from NCBI, *Trypanosoma brucei* (JX910378, JX910373,JN673391,FJ712717,AF306777,AF306774,AF306771andAB742530), *Trypanosoma vivax* (JN673394, KC196703 and TVU22316), *Trypanosoma congolense* (JN673389, TCU22319, TCU22318, TCU22317 and TCU22315), *Trypanosoma simiae* (JN673387 and TSU22320), *Trypanosoma godfreyi* (JN673385) *Trypanosoma evansi* (D89527), *Trypanosoma otospermophili* (AB175625), and *Trypanosoma grosi* (AB175624). They were aligned in Geneious 9.1.5 software (Biomatters Ltd, Auckland, New Zealand) using MAFFT multiple aligner with default settings and ITS1 region identified by comparing annotations and terminal regions of 18s and 1.5s rRNA regions. Primers flanking the ITS1 region were designed and manual sequence editing of the primers was done to improve the range of Trypanosome species and subgroups.

The new primers named AITSF and AITSR were analyzed and the expected amplicon sizes compared with primer pairs of three widely used primers for ITS1 region; CF/BR [18] and ITS1/ITS2 [22] for specificity range with a computer-based *in silico* PCR analysis by Simulate_PCR [23] using the NCBI *nt* database to deduce the scope of Trypanosome species and subgroups detection and the expected lengths of amplicons (S1 Table). Simulate_PCR uses BLAST to search amplicons from a specified database wherein we used a local *nt* database downloaded from NCBI: ftp://ftp.ncbi.nlm.nih.gov/blast/db/ on 3^rd^ December 2017. The new primers were tested using two positive controls; stock DNA of known Trypanosome species and tsetse-derived DNA samples previously confirmed as Trypanosome positive and compared the band sizes with Simulate_PCR results. Simulate_PCR was run using the command;

*simulate_PCR –db <path/to/database> -primers <path/to/primers.fasta> –minlen 100 –maxlen 750 -mm 1 –num_threads 8 –max_target_seq 10000 –genes 1 –extract_amp 1*

We tested the sensitivity of AITSF/AITSR primers against the CF/BR primers to determine their specificity. The ITS1 sequences; *T. brucei* (AF306774), *T. simiae* (JN673387), *T. vivax* (KM391828), *T. congolense* (U22317) and *T. godfreyi* (JN673384) were downloaded from NCBI, synthesized and each insert cloned into a pGEMT-easy vector. Solutions with increasing insert copies were prepared by serial dilution and used as templates for PCR reaction using either AITSF/AITSR or CF/BR primers. Results were analyzed on 5% Agarose gel.

### Paired-end library preparation

A two-step PCR protocol for the library preparation was applied in the multiplex PCR analysis. We used the newly designed AITSF and AITSR primers ligated to Illumina adapter sequences (Table 1).

**Table 1.**
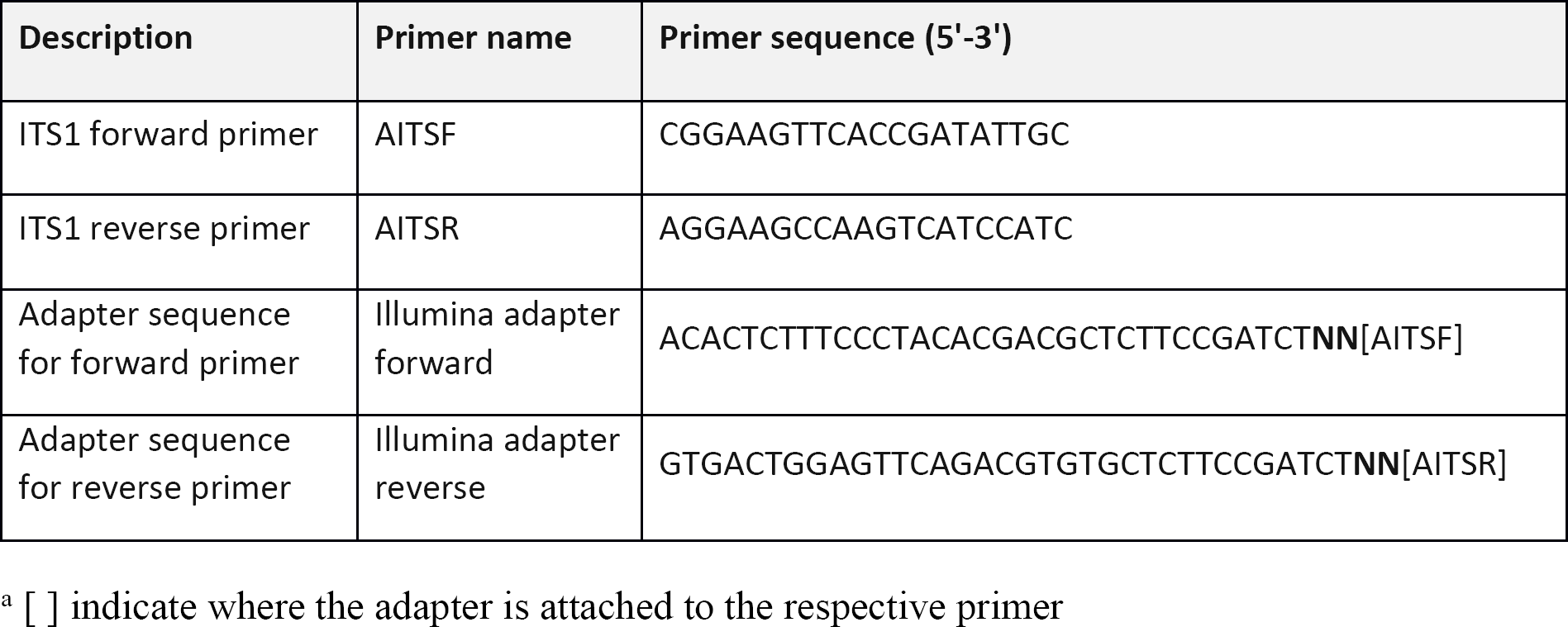
Primers used in this study.

ITS1 PCR was done in duplicate for Rufunsa samples to validate Trypanosome detection results. We also included positive template controls comprising. *T. b gambiense, T. b rhodesiense*, and *T. congolense* DNA. An artificial Trypanosome DNA mixture was included to mimic a mixed infection control. It comprised artificially mixed *T.b. gambiense and T. congolense* DNA mixed in equal proportions. The controls were processed the same as samples from PCR to sequencing. The first PCR reaction used which were ordered in adapter ligated forms where Illumina adapter sequences were added to the 5’ end of each primer (Table1). Sequencing libraries were prepared according to the Illumina MiSeq system instructions [24] substituting with the respective primers. The 20 µL primary reactions contained 0.5 µL of 10 µM each of the forward and reverse primers, 10 µL of 2X Ampdirect® Plus buffer, 0.16 µL of 5 U/μL Taq polymerase (Kapa Biosystems, Boston, USA), 0.4 µL DMSO, and 1 µL extracted DNA as a template. The temperature and cycling profile included incubation at 95°C for 10 min, followed by 37 cycles as follows: 95°C for 30 sec, annealing at 60 °C (for both ITS1 and blood meal primer sets) for 1 min, 72°C for 2 min, final extension at 72°C for 10 min. The 20 µL second PCR reactions contained 1 µL of 10 µM Illumina dual-index primer mix (i5 and i7 primers), 1.2 µL of 25 mM MgCl2, 0.4 µL of 10 mM each of the dNTPs, 0.1 µL of 5 U/μL Taq polymerase, 4 µL 5X buffer, and 2 µL of the 1/60 diluted primary PCR product as template. The temperature and cycling profile included incubation at 95°C for 3 min, followed by 11 cycles as follows: 95°C for 30 sec, 61°C for 1 min, 72°C for 2 min, final extension at 72°C for 10 min. A negative template control was included in each set of PCR reactions. To enable sequencing of all amplicons in this study in one run, we used different sets of dual index primers for each sample in the second PCR reactions.

### Library sequencing

The barcoded second PCR products were analyzed in 1.5% agarose gel. Equal volumes of each sample were pooled into one library. The library pool was purified using the Wizard SV Gel and PCR Clean-Up System (Promega, Madison, WI, USA) by cutting out bands of interest to separate them from primer dimers and post PCR reagents. Quantification of each of the library was done using a Qubit dsDNA HS assay kit and a Qubit fluorometer (ThermoFisher Scientific, Waltham, MA, USA). The concentration of the library was then adjusted to a final concentration of 4 nM using nuclease-free water and applied to the MiSeq platform (Illumina, San Diego, CA, USA). Sequencing was performed using a MiSeq Reagent Kit for 300 base pairs, paired-end (Illumina, San Diego, CA, USA) and a 20% PhiX DNA spike-in control added to improve the data quality of low diversity samples, such as single PCR amplicons. All controls were also included in the sequencing library.

Data obtained from this study is available at SRA database under the SRA accession number SRP159480 (https://www.ncbi.nlm.nih.gov/sra/SRP159480).

### Bioinformatics

The analysis followed a workflow (Fig 2) comprising the AMPtk pipeline coupled with taxonomic identification by BLAST. All commands for analysis were run as a custom script (S1 Text). Briefly, reads were processed using the AMPtk pipeline by; 1) Trimming primers, removal of sequences less than 100 b.p, and merging pair-end reads. Merging parameters were customized by editing the AMPtk file amptklib.py with the USEARCH options; *fastq_pctid* set to 80, (minimum %id of alignment), *minhsp* set to 8, and *fastq_maxdiffs* set 10 to limit the number of mismatches in the alignment to 10. 2) Clustering; the DADA2 denoising algorithm option was called using the *amptk dada2* command. This algorithm provides a clustering independent method that attempts to “correct” or “denoise” each sequence to a corrected sequence using statistical modeling of sequencing errors. AMPtk implements a modified DADA2 algorithm that produces both the standard “inferred sequences” referred to as amplicon sequence variants (ASVs) output and also clusters the ASVs into biologically relevant OTUs using the UCLUST algorithm. 3) Downstream processing of ASVs where ASV table filtering was done to correct for index-bleed where a small percentage of reads bleed into other samples. This was done by the *amptk filter* command using 0.005, the default index-bleed percentage. 4) An additional post-clustering ASV table filtering step was done using the *amptk lulu* command. LULU is an algorithm for removing erroneous molecular ASVs from community data derived by high-throughput sequencing of amplified marker genes [25]. LULU identifies errors by combining sequence similarity and co-occurrence patterns yielding reliable biodiversity estimates. 5) Taxonomy was assigned to the final ASV (OTU) table. ASV taxonomic identification (in this study) was done by BLAST (v2.6.0) [26] remotely. The BLAST output file was parsed and edited to match the taxonomy header formatting specified in the AMPtk manual and subsequently used for generating a taxonomy labeled ASV table.

**Fig 2.**
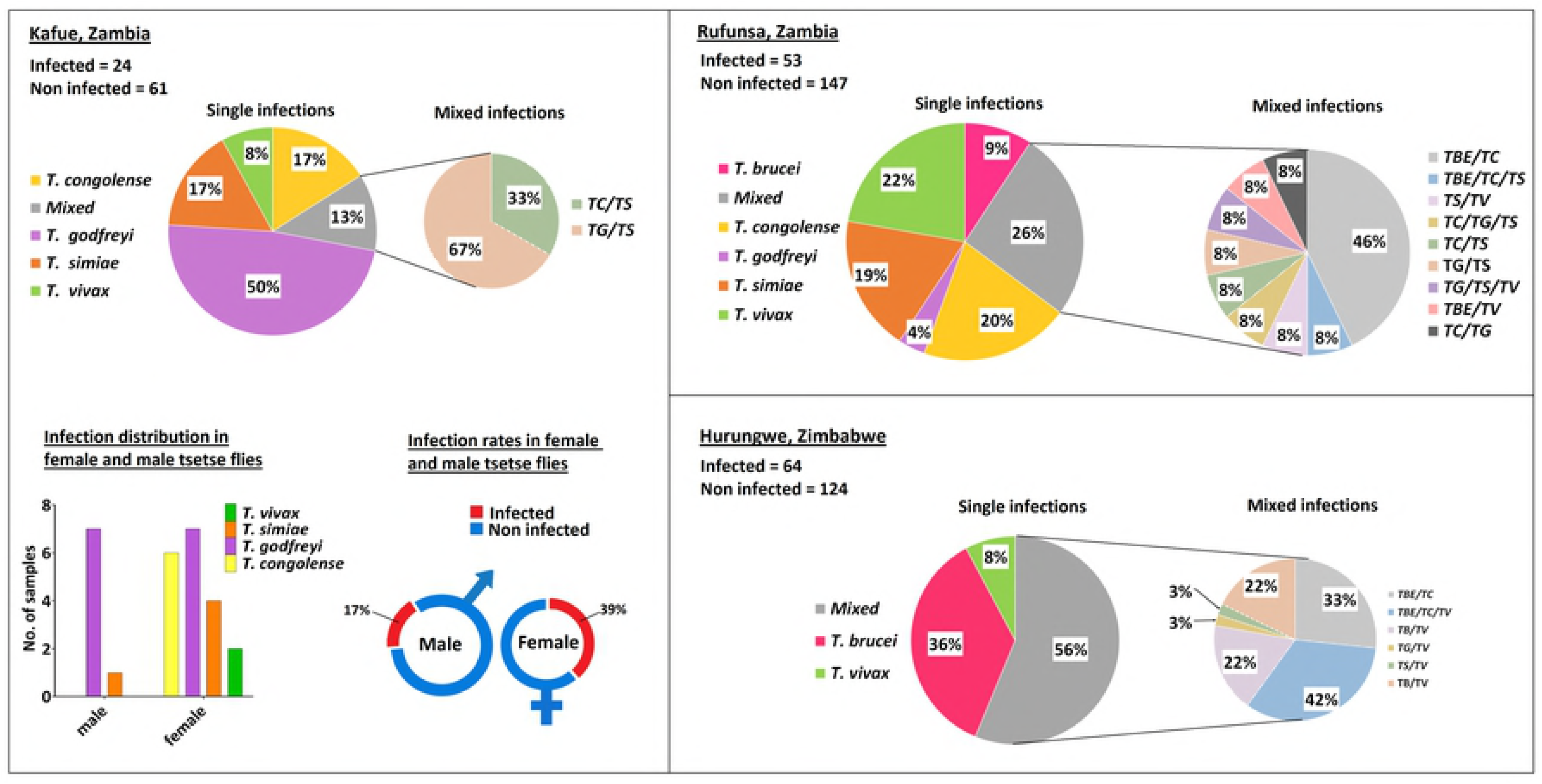
Workflow for read analysis using AMPtk pipeline.

To check the accuracy of the ASVs generated by the Amptk pipeline, we simulated FASTQ files generated *in silico* from downloaded sequences used in a previous study [11]. This was done by running *ArtificialFastqGenerator* [27], to generate paired-end FASTQ files with 1000 reads per sequence. Real quality scores and simulation of sequencing errors was achieved by using a pair of FASTQ files from sequencing output of the samples. Amptk pipeline was then run on the generated reads. The resultant ASVs were allocated taxonomic identity at species level by BLAST and then compared to the species identity of parent sequences. All the software used in data analysis are free under open access licenses.

### Phylogenetic and statistical analysis

A phylogenetic tree was created from the alignment generated from ASVs obtained after analysis. Alignments were made with MAFFT [28] using the *mafft-xinsi* option (allowing for prediction of RNA secondary structure and build a multi-structural alignment) with 1,000 maximum iterations, leaving gappy regions and using kimura 1 option for score matrix. Maximum likelihood phylogenetic trees were built with RAxML 8.0.26 using the ‘GTRCATI’ model and default parameters with 10,000 bootstraps. The tree was visualized and annotated using iTOL (version 4) [29]. Statistical analysis and graphing of data were carried out in GraphPad Prism version 6.01 for Windows, GraphPad Software, San Diego California USA, www.graphpad.com.

## Results

### Improved primers

We evaluated newly designed primers (AITSF/AITSR) and compared their sensitivity to conventionally used ITS1 primers; CF/BR primers [18]. PCR performed on pGEMT-easy plasmid DNA with ITS1 inserts from different Trypanosome species at different dilutions showed that the new primers were slightly more sensitive (S1 Fig). PCR done using AITSF/AITSR primers were able to detect as little as 10^2^ *T. godfreyi* inserts, 10^3^ *T*. *simiae, T. vivax* and *T*. *congolense* inserts and up to 10^4^ *T*. *brucei* inserts while CF/BR primers detected 10^3^ *T*. *godfreyi* and *T*. *vivax* inserts and 10^4^ *T*. *simiae* and *T*. *congolense* inserts.

### Read data and replicate analysis

Reads generated from amplicon sequencing were of relatively good quality. Apart from those from Zimbabwe, more than 90% of the reads passed quality filtering in all samples (Table 2). The no. of ASVs generated in replicate runs was slightly different indicating slightly different detection sensitivities in the replicate PCR runs. Only the forward read was retained for downstream analysis in reads that did not merge due to either amplicon being longer than 600 b.p or due to low-quality bases in the overlap bases. This did not affect the final identification of reads as shown by the simulated data results described later. We analyzed the Rufunsa samples in replicate and compared the results. Both replicates had similar results in regard to individual Trypanosome species detection per sample seen in the gel image analysis (Fig 3A) as well as amplicon read analysis (Fig 3B). The outcome of detection for each of the Trypanosome species and sub-groups in replicate runs was comparable and the Fischer’s exact test confirmed that there was no significant difference (*P<0.05*) in the number of positive detections in replicate runs (S2 Table).

**Table 2.** Read data of all samples analyzed.

| Source of sample | No. of samples | Total no. of reads | Reads after pre-processing (% of total) | Raw ASVs | OTUs (97% clustering of ASVs) | ASVs post-filtering |
| --- | --- | --- | --- | --- | --- | --- |
| Rufunsa Run A | 200 | 916,055 | 897,598 (99.8%) | 269 | 89 | 174 |
| Rufunsa Run B | 200 | 1,289,667 | 1,248,934 (94.8%) | 320 | 95 | 232 |
| Kafue | 85 | 483,589 | 454,799 (91.4%) | 131 | 48 | 56 |
| Hurungwe | 188 | 29,798 | 11,247 (79.5%) | 137 | 63 | 116 |

ASVs generated were filtered to remove underrepresented and/or artifact ASVs from the final taxonomy table.

**Fig 3.**
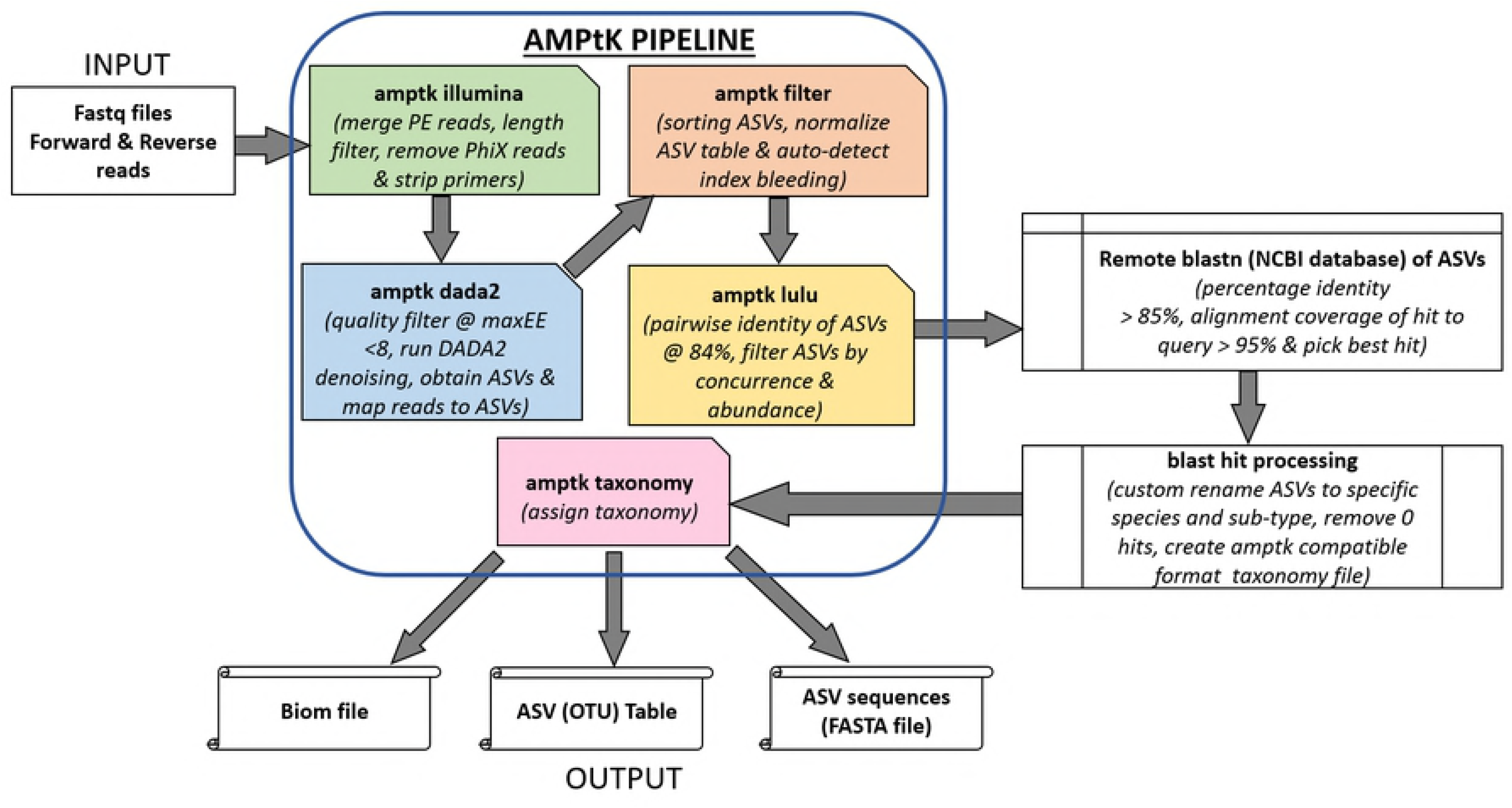
Representative replicate analysis results. (A) Gel analysis of Rufunsa samples done in replicate showing matching bands per sample. (B) Amplicon sequence analysis of the same samples in A) showing number of reads detected per species in each sample.

### Pipeline validation and accuracy of detection

Simulation of data generated from Trypanosome sequences downloaded from NCBI and analyzed using the AMPtk (amplicon toolkit) pipeline (version 1.2.4) (https://github.com/nextgenusfs/amptk) showed that amplicon sequence variants (ASVs) generated by the pipeline as primary units of representing sequence diversity, were more accurate in correctly inferring the diversity sequences compared to operational taxonomic units (OTUs) derived from clustering sequences at 97% identity (S3 Table). The specificity and precision of distinguishing between individual sequences of the same Trypanosome species are reflected by the number of ASVs or OTUs representing each of the different species. For example, only one OTU was generated for all three *Trypanosoma theileri* sequences, and three OTUs were generated for seven *Trypanosoma simiae* sequences, while the number of ASVs generated in each case represented each sequence accurately. The simulated data results indicated that read analysis using the AMPtk pipeline and ASVs instead of OTUs was suitable for sensitive identification of Trypanosome reads.

### Amplicon sequencing improves the sensitivity of detection and reveals errors of detection in conventional ITS1 PCR-gel analysis

By comparing gel images after PCR and sequence data, it was observed that the sensitivity of detection of Trypanosome DNA was increased after sequencing. Samples with bands that were barely visible after the 1^st^ PCR became visible after the 2^nd^ PCR and were confirmed as positive after sequencing (Fig 4A). It was also observed that some *T. godfreyi* and *T. vivax* amplicon bands were of a relatively similar size and it was difficult to distinguish the two by gel analysis alone (Fig 4B). Mixed and single infections with multiple and single bands respectively were observed and confirmed by amplicon sequence analysis. Results for the second PCR using dual-index primers showed consistency with those of the first PCR. There were no bands visible outside the expected range indicating the absence of non-specific amplification in both PCR steps. The 1^st^ PCR amplicons were slightly longer than expected sizes due to the adapter sequences (approx. 80 bp) added to the primer, therefore the bands observed corresponded to *T. congolense (Kilifi/Forest and Savannah);* 650-800 b.p, *T. brucei;* 520-540 bp, *T. simiae;* 440-500 bp, *T. godfreyi;* 320-400 bp, and *T. vivax;* 290-400 bp.

**Fig 4.**
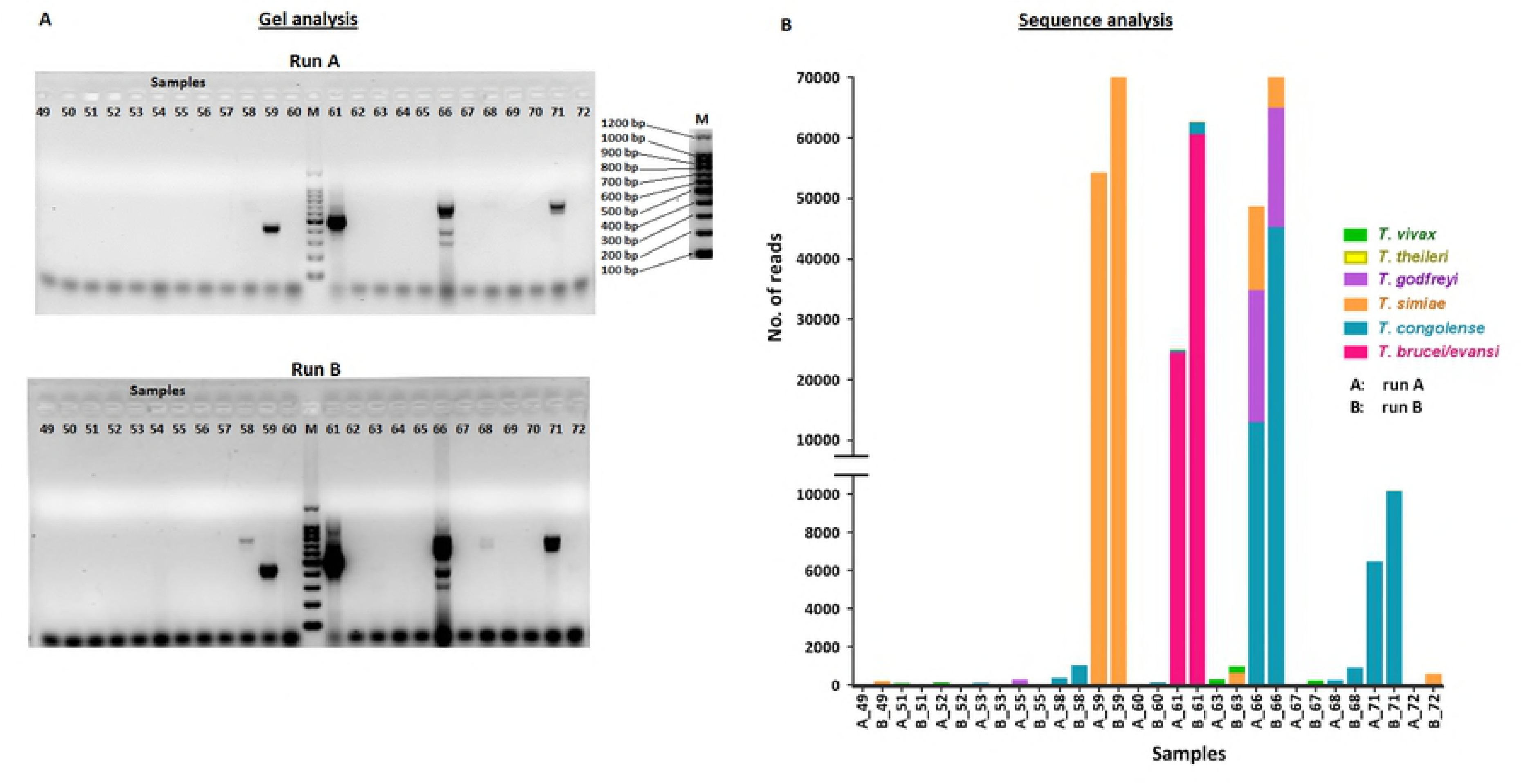
Representative gel and sequence analysis results. (A) Arrows showing bands are not visible after the 1st PCR become visible after 2nd PCR. (B) By gel analysis, amplicon bands of samples 5, 7 and 10 are indistinguishable by size and are deemed to be all *T. godfreyi* while sequencing reveals that the amplicon of sample 10 is, in fact, *T. vivax*. Positive controls comprise; Tbg (*T. brucei gambiense*), Tbr (*T. brucei rhodesiense*), Tb/Tc (an artificial mixture of equal amounts of *T. brucei gambiense* and *T. congolense* DNA).

### Trypanosome ITS1 sequences can be used to distinguish between Trypanosome species and subgroups

The accuracy in distinguishing between Trypanosome species and subgroups was analyzed by phylogenetic analysis of ASV sequences and their species identity allocated by BLAST. ASVs were named after the accession number of their respective top hit BLAST subject sequence and area of collection of the sample they originated from. Phylogenetic analysis of all ASVs obtained from this study showed that ASVs named after same Trypanosome species clustered together regardless of sample collection location. Sub-clustering into different subgroups of the same species was also observed (Fig 5). The *Nannomonas* subgenus showed the highest diversity of sub-clustering where *T. simiae* clustered into two main subgroups; *T. simiae* and *T. simiae Tsavo*. Two *T. simiae* Tsavo II ASVs from Kafue, with 91% and 97% identity to *T. congolense* Tsavo (Accession number U22318) recently reviewed and classified as *T. simiae* Tsavo [30,31] clustered distinctly from the rest of the *T. simiae* Tsavo I ASVs. *T. congolense* ASVs showed the highest diversity and clustered into three main subgroups; Kilifi, Riverine/Forest, and Savannah. *T. congolense* Savannah represented the most diversity in all the ASVs analyzed from all the samples. *T. congolense* Kilifi clustered separately and far from *T. congolense* Savannah and Riverine/Forest subgroups. *T. godfreyi* showed sub-clustering into two main sub-groups while *T. vivax* (belonging to the *Dutonella* subgenus) also clustered into two sub-groups. The *Trypanozoon* subgenus (*T. brucei/T. evansi*) did not show any distinct sub-clustering.

**Fig 5.**
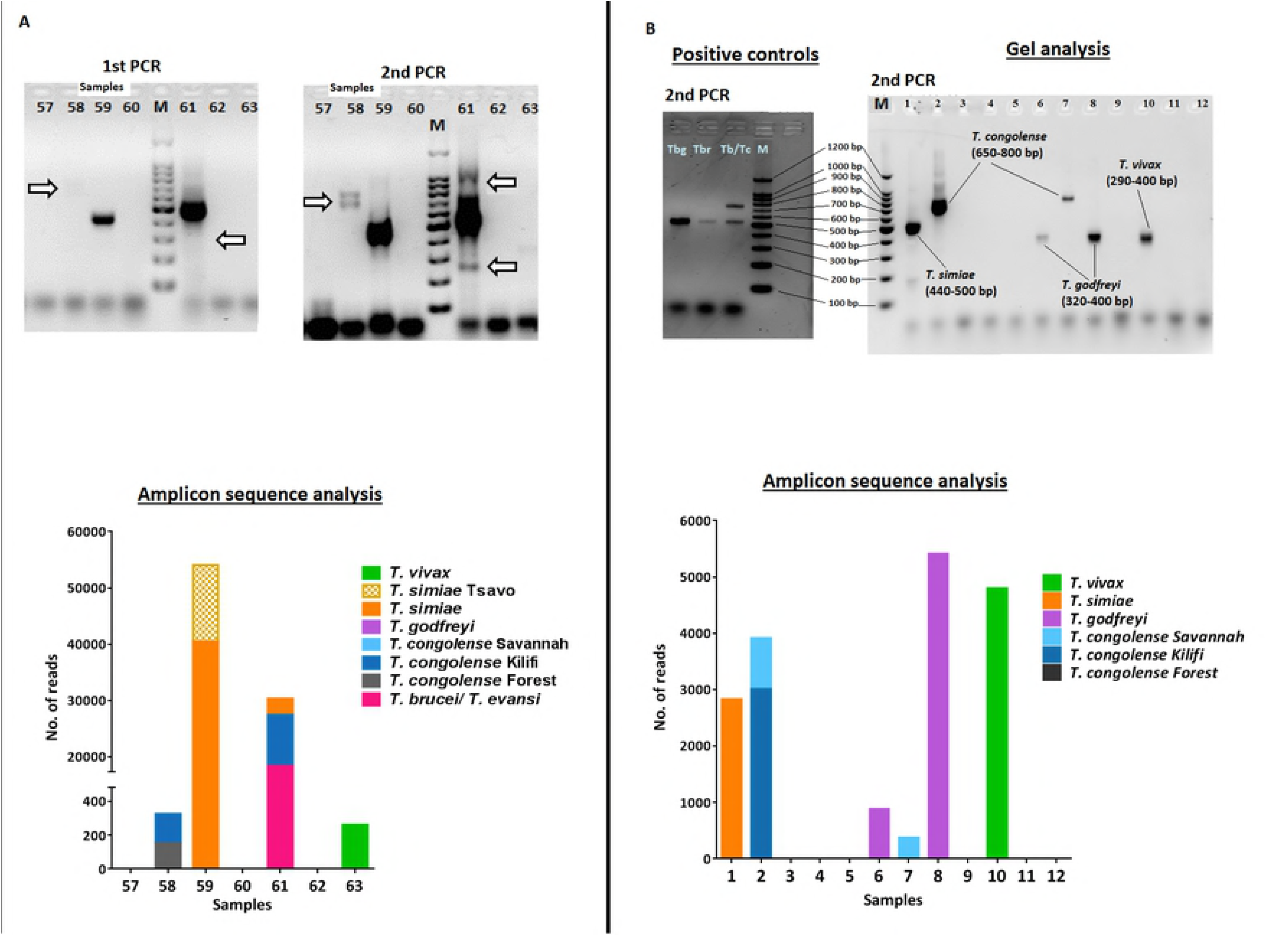
Phylogenetic tree of unique ASVs generated from amplicon sequence data. A *Bodo caudatus* ITS1 sequence was included as outgroup. Individual Trypanosome species and subgroups cluster into distinct clades. ASV are named after their respective blast best hit matches.

### Prevalence and distribution of Trypanosome species

The prevalence of Trypanosome infection in the Rufunsa area, Zambia, was 25.6%, that of in the Kafue area, also Zambia, 28.2%, while that of the Hurungwe area, Zimbabwe, was 47.3%. Flies caught in Rufunsa had the highest prevalence of *T. congolense* while those from Kafue had the highest prevalence of *T. godfreyi* (Table 3). The highest prevalence of *T. brucei/T. evansi* was recorded in flies caught in Hurungwe. We did not detect any *T. brucei*/*T. evansi* from flies collected in Kafue. Mixed infections were predominant in flies caught in Rufunsa and Hurungwe while flies caught in Kafue were predominantly infected with *T. godfreyi* (Fig 6). Only tsetse flies from the Kafue region were sorted by sex during collection and we observed that the infection rate in female flies (38.6%) was more than twice that of male flies (17.1%). Additionally, we did not detect *T. congolense* and *T. vivax* infections in male flies. Flies caught in Hurungwe did not have single infections with *T. congolense* or *T. godfreyi.*

**Table 3.** Prevalence of Trypanosome species infection in caught tsetse flies.

| Trypanosome species | Rufunsa (n=200) | Kafue (n=85) | Hurungwe (n=188) |
| --- | --- | --- | --- |
| ***Trypanozoon*** | 6.0%<br>(3.5% - 10.2%) | 0.00%<br>(0% - 4.3%) | 45.7%<br>(38.8% - 52.9%) |
| ***T. congolense* Forest** | 4.5%<br>(2.4% - 8.3%) | 1.2%<br>(0.2% - 6.4%) | 0.0%<br>(0% - 2.0%) |
| ***T. congolense* Kilifi** | 7.5%<br>(4.6% - 12.0%) | 2.4%<br>(0.7% - 8.2%) | 4.8%<br>(2.5% - 8.9%) |
| ***T. congolense* Savannah** | 7.5%<br>(4.6% - 12.0%) | 4.7%<br>(1.9% - 11.5%) | 39.9%<br>(33.2% - 47.0%) |
| ***T. godfreyi*** | 3.0%<br>(1.4% - 6.4%) | 16.5%<br>(10.1% - 25.8%) | 3.7%<br>(1.8% - 7.5%) |
| ***T. simiae*** | 6.0%<br>(3.5% - 10.2%) | 5.9%<br>(2.5% - 13.0%) | 1.1%<br>(0.3% - 3.8%) |
| ***T. simiae* Tsavo** | 8.7%<br>(4.5% - 16.2%) | 2.4%<br>(0.7% - 8.2%) | 0.0%<br>(0% - 2.0%) |
| ***T. vivax*** | 7.5%<br>(4.6% - 12.0%) | 2.4%<br>(0.7% - 8.2%) | 29.2%<br>(23.2% - 36.1%) |
| **Trypanosoma**<br>**(overall prevalence)** | 26.5%<br>(20.9% - 33.0%) | 28.2%<br>(19.8% - 38.6%) | 47.3%<br>(40.3% - 54.5%) |

Confidence levels at 95% for apparent prevalence (Wilson) are shown in brackets.

**Fig 6.**
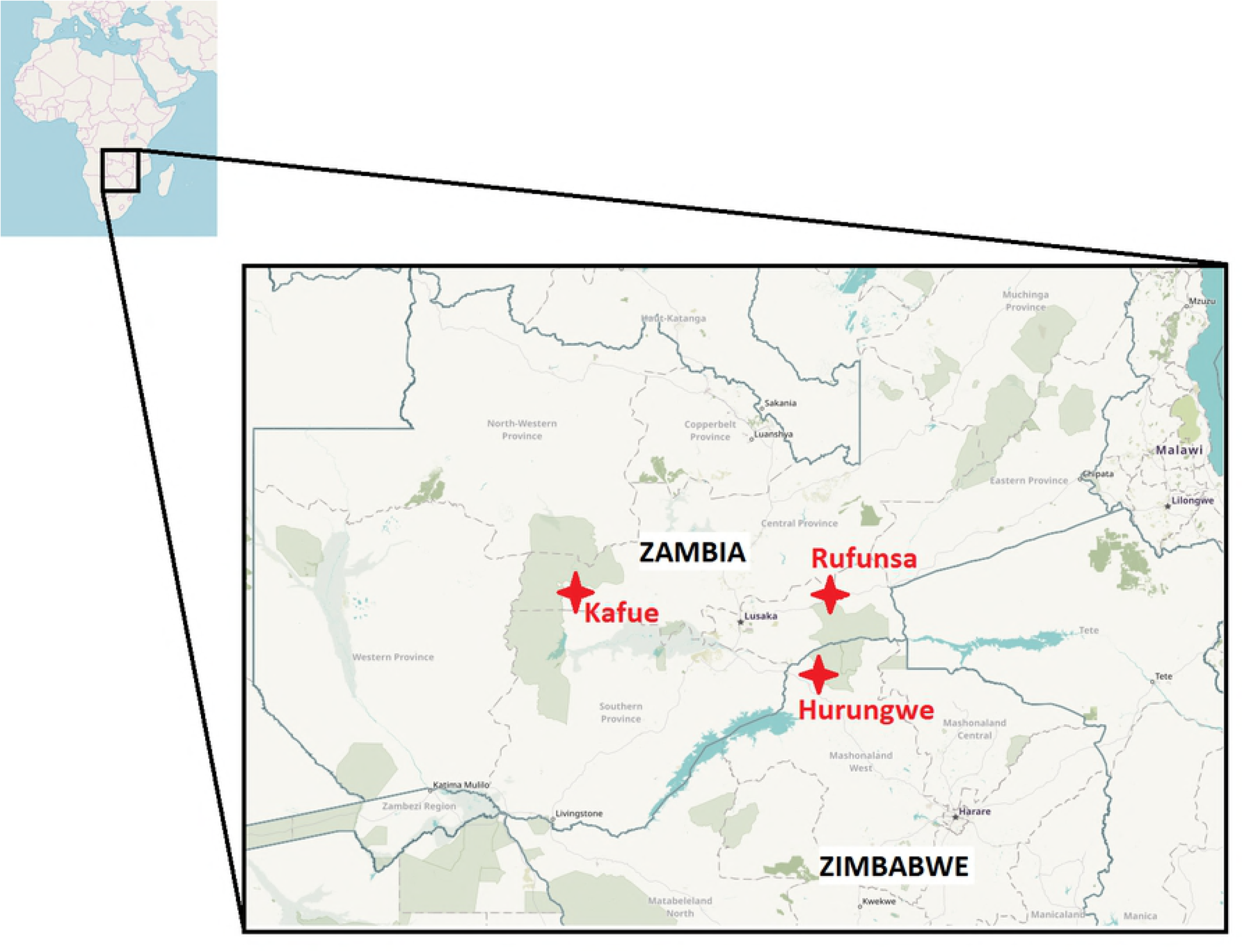
The distribution of Trypanosome species amongst infected tsetse flies. TBE = *T. brucei/T. evansi*, TV = *T. vivax*, TS = *T. simiae*, TG = *T. godfreyi*, and TC = *T. congolense*.

## Discussion

This study reports a new and versatile approach for detection of Trypanosome DNA in samples with high sensitivity and precision than conventional PCR-gel approach. We have established that conventional ITS PCR gel analysis is not an accurate way of determining the prevalence of Trypanosome species infections since identification of species by band size is inaccurate and may lead to misidentification of some Trypanosome species. Apart from the *Trypanozoon* group (*T. brucei and T. evansi*) which are extensively similar at the genome level [32], our new approach is sensitive at the subgroup level and has a high capacity to process large amounts of samples in one run (approximately a 700 samples mixed library) owing to the high repertoire of Illumina dual indexing primers. As part of this work, we have also developed new primers that are more sensitive than conventional primers and cover a wider range of the *Trypanosoma* genus. With our approach, it is now possible to identify species and subgroups of Trypanosomes by sequence analysis on individual samples as opposed to pooled samples for a large dataset which allows for the detection of new isolates. It is also possible to make a better inference of the Trypanosome species circulating in an area. This approach is a practical and, with the decreasing cost of next-generation sequencing, cost-effective way to monitor large field samples of all kinds. They can, therefore, be utilized in a wide range of samples from vectors and hosts and the analysis of new Trypanosome species.

The results obtained in this study indicate that *T. vivax* and *T. godfreyi* have very similarly sized ITS1 amplicons making it difficult to identify one from the other based solely on gel band sizes. Sequencing and clustering of the reads effectively address this issue.

Phylogenetic analysis shows several interesting population substructures in the cases of *T. simiae* and *T. congolense*. Within the *T. congolense* clade, Savannah and Riverine/Forest subgroups show more sequence similarity while the Kilifi type shows more divergence. This agrees with a previous study that found *T. congolense* Savannah and Riverine/Forest had 71% similarity in satellite DNA sequence [33] and that the Kilifi subgroup was as divergent from other *T. congolense* subgroups [34]. The clustering of *T. congolense* Kilifi close to *T. simiae* species than other *T. congolense* subgroups is quite interesting in that an earlier study had identified a new *T. congolense* Tsavo strain (Accession number U22318) [35] which has been classified as *T. simiae* Tsavo [36]. We identified two ASVs from Kafue area (classified as *T. simiae* Tsavo II in this study) that had 91% and 97% identity to the U22318 *T. congolense* Tsavo sequence and that clustered with *T. simiae* Tsavo rather than other *T. congolense* species sequences supporting the T. simiae Tsavo classification. However, they cluster separately from the other *T. simiae Tsavo* ASVs, suggesting that they may have a divergent genotype. Perhaps there is a complex relationship between *T. congolense* and *T. simiae* species yet to be identified.

Prevalence of Trypanosome differed between the sampled areas with single and mixed infection being detected in flies caught agreeing with previous studies [20,37,38]. This may be an important factor in the exchange of information between species. We also observed that the infection rate of female tsetse flies was twice that of male flies. This result is in contrast with experimental studies using laboratory maintained tsetse flies that found males being more susceptible than females [39–41].

To conclude, our results imply that with the new primers, it is possible to detect and distinguish between different Trypanosome species and subgroups accurately and therefore infer prevalence of infection more precisely using a single test without having to undertake satellite DNA analysis that requires species-specific primers. This is made possible by deep sequencing which enables resolution at a single nucleotide level. This high resolution at sub-cluster level utilizing only the ITS1 region has not been shown before thus a practical and sensitive barcoding of African trypanosomes. Using our approach, it is thus possible to distinguish *T. godfreyi* from *T. vivax*, as well as highlight finer subpopulation structures within the *T. simiae* and *T. congolense* clades that raise interesting questions regarding their classification. It is highly likely that there are genomic and taxonomic differences between *T. vivax, T. godfreyi* and *T. congolense* sub-groups that need to be studied. This could provide answers on the evolution of Trypanosomes. WhatcontributiondotheseTrypanosomesubgroupsmaketolivestockdisease?Arethese genotypes responsible for assumed “strain” differences in drug response? Can these new genotypes be correlated with the old morphological criteria and species designations? Do these “strains” have the potential of evolving to new subgroups that could pose new risks? There is a need for more studies to catch up with the molecular taxonomy to answer these questions.

## Acknowledgements

We wish to acknowledge Mr. Lambert Gwenhure for his assistance in obtaining samples from Zimbabwe as well as staff of Hokudai Center for Zoonosis Control in Zambia (HCZCZ) for their help during collection of samples in Zambia.

---

**S1 Fig. Sensitivity of AITSF/AITR primers compared to CF/BR primers in the detection of Trypanosome ITS1 inserts cloned in the pGEMT-easy vector.**

**S1 Text. Script with all commands used to run the AMPtk pipeline.S1 Table. Amplicon sizes of new primers (ATSF/AITSR) compared to other primers (CF/BR and ITS1/ITS2) obtained by simulated PCR.**

**S2 Table. Statistical analysis of detection of individual Trypanosome species in replicate runs.**

**S3 Table: Matrix comparison of ASVs and OTUs from simulated data.**

